# Receptor tyrosine kinases regulate signal transduction through a liquid–liquid phase separated state

**DOI:** 10.1101/783720

**Authors:** Chi-Chuan Lin, Kin Man Suen, Polly-Anne Jeffrey, Lukasz Wieteska, Amy Stainthorp, Caroline Seiler, Hans Koss, Carmen Molina-París, Eric Miska, Zamal Ahmed, John E Ladbury

## Abstract

Receptor tyrosine kinases (RTKs), the largest class of transmembrane cell surface receptors, initiate signalling pathways which regulate diverse cellular processes. On activation these receptors rapidly recruit multiple downstream effector proteins to moderate affinity tyrosyl phosphate (pY) binding sites. However, the mechanism for expedient downstream effector protein recruitment via random molecular diffusion through the cytoplasm is not fully understood. One way in which the probabilistic outcome associated with random diffusion could be alleviated is through localized accumulation of high effective concentrations of signalling proteins in discrete pools in the cell (*1*). The inclusion of interacting proteins into liquid-liquid phase-separated (LLPS), membraneless protein droplets maintains functionally relevant proteins at high concentrations in a liquid phase at the required point of action, enhancing equilibrium binding and enzyme activity (*2–6*). These LLPS states have been associated with a wide range of cellular functions including regulation of signalling through, for example, nephrin (*7, 8*), the T-cell receptor (*9*), mTOR (*10*), and Sos-Ras (*11*), however, whether LLPS extends to RTK-mediated signal transduction has not been investigated. Here, we show that an RTK, fibroblast growth factor receptor 2 (FGFR2), forms a signalling competent LLPS state with two downstream effectors, a tandem Src homology 2 (SH2) domain-containing protein tyrosine phosphatase 2 (Shp2), and 1-phosphatidylinositol 4,5-bisphosphate phosphodiesterase gamma 1 (Plcγ1). We show that these proteins assemble into a ternary complex which exploits LLPS condensation to simultaneously modulate kinase, phosphatase and phospholipase activities. Therefore, LLPS formation ensures that the requirement for prolonged, high-fidelity signalling is achieved. Additional RTKs also form LLPS with their downstream effectors, suggesting that formation of biological condensates is a key organising principle of RTK-mediated signalling, with broad implications for further mechanistic studies as well as therapeutic intervention.

---

The existence of LLPS properties was investigated for a subset of phosphorylated RTKs (pEGFR, pHer2, pHer4, pFGFR1, pFGFR2, pVEGFR1 and pVEGFR2) and the downstream effector proteins, phosphatase Shp2 (the inactive C459S mutant, Shp2_C459S_ (*12*)) and adaptor Shc (Fig. 1A). Clearly visible droplets are apparent including both EGFR- and FGFR-family RTKs with Shp2_C459S_, whilst Shc phase separates with all of the RTKs tested. Although, the formation of RTK-mediated droplets was observed across the panel of RTKs, we selected the phase separation in the presence of FGFR2 and Shp2 for further investigation. Compared to the basal state, stimulation of FGFR2 results in coalescence of micrometer-sized clusters at the plasma membrane of complexes containing Shp2 (Fig. 1B left panel). Another well-studied substrate protein of pFGFR2, Plcγ1 (*13*), was also seen to condense into droplets on stimulation of cells (Fig. 1B middle panel), whilst no such clusters were observed in the absence of the receptor (Fig. 1B right panel). To explore the mechanism of formation, and functional consequences of FGFR2-containing microclusters, we reconstituted the condensed state from purified components. Sub-micrometer-sized droplets formed upon addition of Shp2 and Plcγ1 to pFGFR2 cytoplasmic domain (pFGFR2_Cyto_) in the presence of ATP/Mg^2+^. Replacing Shp2 with the inactive mutant Shp2_C459S_ resulted in enhanced droplet formation, suggesting that phase separation is dependent on prolonged phosphorylation (Fig. 1C). Further examination revealed that droplet formation was absent with the individual proteins, but was apparent with the pFGFR2_Cyto_/Shp2_C459S_ and Shp2_C459S_/pPlcγ1 pairwise combinations (fig. S1, A and B). The lack of droplet formation between pFGFR2 and pPlcγ1 is consistent with the reported abrogation of interaction upon Y783 Plcγ1 phosphorylation (*14, 15*). To provide robust confirmation of LLPS state formation, we employed three complementary methods that have emerged as a current standard in the field (*16*). As expected for liquid-liquid droplets, addition of 1,6-hexanediol led to complete dispersal of the condensates (fig. S1A(vi)), as did the addition of sodium chloride (NaCl) (fig. S1A (vii, viii)). Additionally, the fluorescence recovery after photo-bleaching (FRAP) is consistent with a fluid state (Fig. 1D). We next examined the nature of the complex(es) formed by the three components of the LLPS state. First, we characterised the pairwise protein interactions involved in mediating the ternary complex formation. We demonstrated that Shp2 and FGFR2 engaged using multivalent, cooperative interactions mediated by the C-terminal SH2 domain of Shp2 (CSH2) bound to phosphorylated Y769 on the C-terminal tail of the receptor (Fig. 2A to C, fig. S2, A to D and fig. S3, A to D) as well as additional non-specific weak interactions between Shp2 NSH2 and secondary pY sites on the kinase domain (Fig. 2A, fig. S3D, and table S1).

**Fig. 1.**
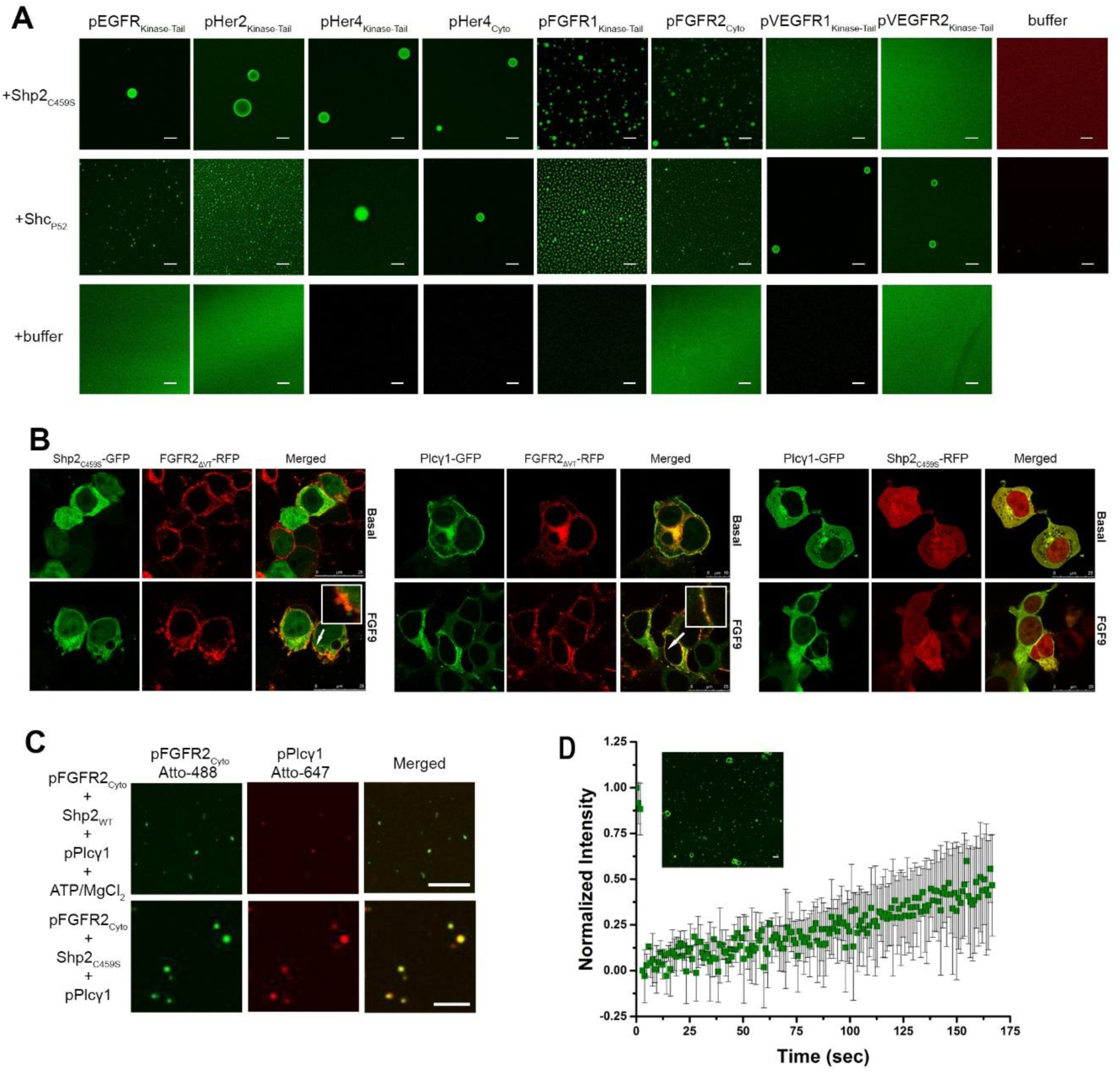
Phosphorylated RTK-mediated condensation of protein complexes. (**A**) Droplet formation of recombinant phosphorylated EGFR, FGFR, and VEGFR (6 µM each, Atto-488 labelled) upon adding 30 µM of Shp2_C459S_ (top panel) or Shc_P52_ (middle panel). Scale bar = 10 μm. (**B**) Cell images showing droplet formation upon FGFR2 expression and activation. FGFR2 has the VT motif deleted to abrogate indirect binding of Shp2 via the adaptor protein Frs2 (FGFR2_ΔVT_^24–26^). Insets: magnification of regions shown by arrow to exemplify phase separated droplets. No evidence of droplet formation in the absence of FGFR2_ΔVT_ (right panel). (**C**) LLPS state observed between Shp2 (60 µM) and pFGFR2_Cyto_ Atto-488 (6.6 µM) and pPlcγ1 Atto-647 (15 µM). Using inactive Shp2_C459S_ mutant promotes the formation of droplets (lower panel). Scale bar = 10 μm. (**D**) FRAP recovery curve for pFGFR2_Cyto_-Atto488 (10 µM) and Shp2_C459S_ (60 µM) with error bars indicating standard error of the mean. Inset - droplets on photobleaching. (n = 3)

**Fig. 2.**
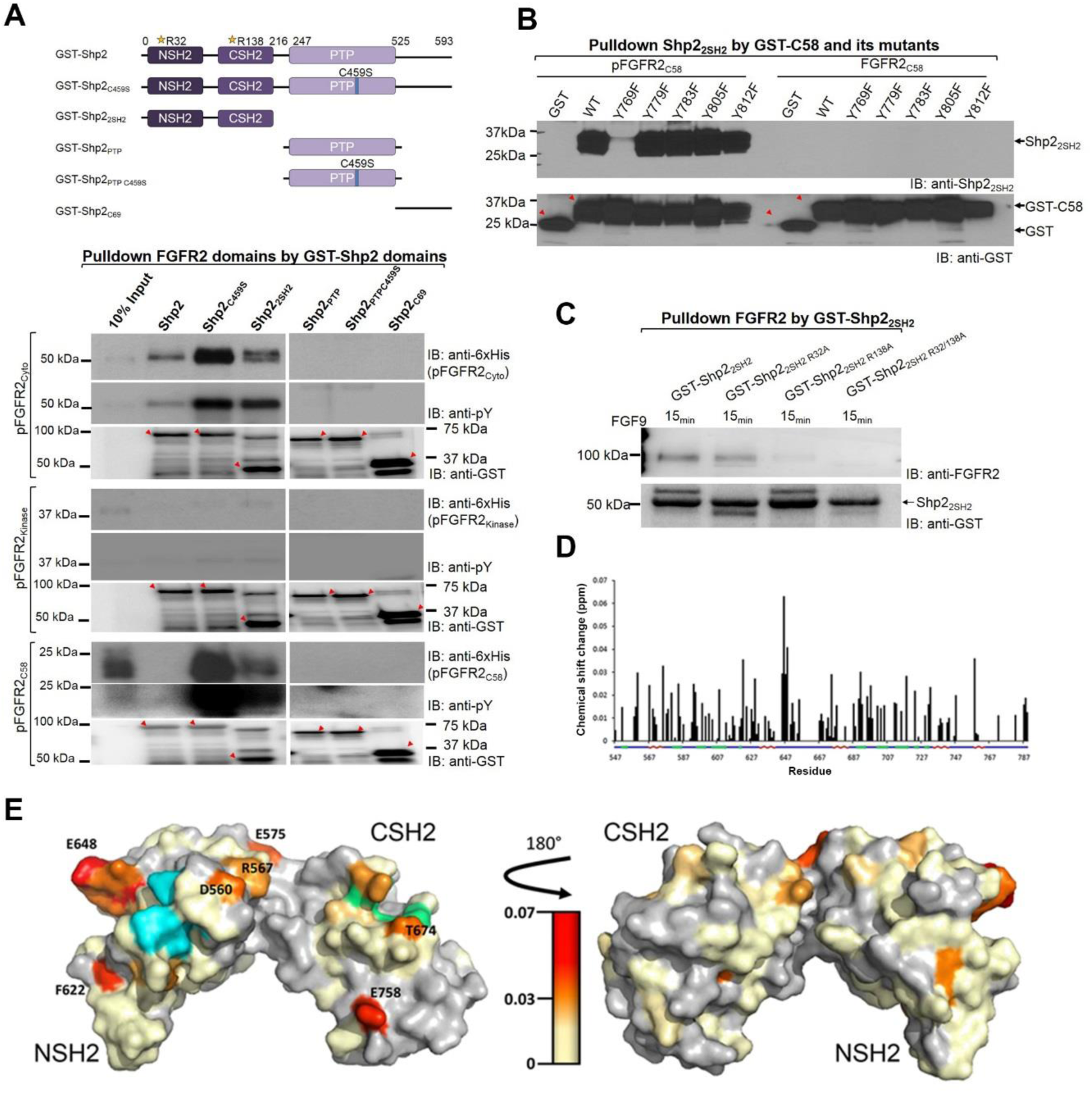
Characterisation of the interactions between droplet-forming components. (**A**) Pull down experiments using different Shp2 constructs (see schematic - the location of the mutated residues R32, R138 and C459 shown). GST-Shp2, GST-Shp2_C459S_ and GST-Shp2_2SH2_ pull down pFGFR2_C58_ and pFGFR2_Cyto_ whilst pFGFR2_kinase_ was pulled down at a significantly lower level. Red arrows - fusion protein loading control. (**B**) The C-terminal 58 residues of FGFR2, GST-FGFR2_C58_ with individual Tyr to Phe substitutions, were phosphorylated and used to pull down Shp2_2SH2_. The Y769F mutation abrogates binding. Red arrows highlight GST and GST-FGFR2_C58_ constructs loading controls. (**C**) GST-Shp2_2SH2_, GST-Shp2_2SH2 R32A_, GST-Shp2_2SH2 R138A_ and GST-Shp2_2SH2 R32/138A_ were used to pull down FGFR2_ΔVT_ from HEK293T cells. Mutation of R138 abrogates binding of FGFR2 confirming the requirement of only the wild type CSH2 domain for binding to receptor. (**D**) Plot of the chemical shift changes (ppm) of the backbone amide peaks of ^1^H, ^15^N-labelled Plcγ1_2SH2_ upon addition of 3 molar equivalent of Shp2_2SH2_. The residue numbers are indicated on the x-axis. The secondary structural elements of Plcγ1_2SH2_ are indicated as follows: β-sheet (green), α-helix (red), unstructured (blue). (**E**) CSP of residues mapped on to the crystal structure of the Plcγ1_2SH2_ (PDB code: 4FBN). The gradient indicates the strength of the perturbation. The pY binding pockets for NSH2 and CSH2 are shown in cyan (R562, R586, S588, E589, T590, and T596) and green (R675, R694, R696 and A703), respectively. Left hand image shows putative binding region (highlighted by increasing CSP). Right hand image shows structure rotated into plane by 180º to show the comparatively negligible CSP on the ‘non-binding’ surface.

Direct binding between Shp2 and Plcγ1 is mediated by the tandem SH2 domains (Shp2_2SH2_ and Plcγ1_2SH2_ respectively) and was found to have a moderate affinity (K_d_ ~ 1.2 ± 0.1 µM; fig. S4A and table S1). Comparison of the nuclear magnetic resonance (NMR) spectra of the apo-Plcγ1_2SH2_, and the Plcγ1_2SH2_-Shp2_2SH2_ complex showed mostly minor, widely distributed changes (fig. S4, B to D) that are consistent with small structural and/or dynamic changes (e.g. associated with domain/subdomain rearrangements). However, a limited number of residues showed pronounced chemical shift perturbations (CSPs) indicative of a specific binding event (Fig. 2D). We mapped observed CSPs onto a structure of Plcγ1_2SH2_ (*15*), and observed that they localised to a potential binding region (Fig. 2E). Binding of Shp2 to this site would occlude the N-terminal pY binding pocket abrogating the binding of Plcγ1 to pY769 on FGFR2. The CSPs of residues connecting the CSH2 domain with the Y783 peptide region on Plcγ1 are known to be sensitive to any direct or allosteric interference due to fast exchange equilibria between bound and unbound states (*17*). The absence of any corresponding CSPs indicates that the previously observed intramolecular interactions of the CSH2 domain on Plcγ1 with the pY783 binding pocket remain largely unperturbed by Shp2 binding, and hence can preserve the active state of the phospholipase. This ensures that when both proteins are present only Shp2 can engage the receptor. The presence of pY783 on pPlcγ1 increases the affinity of the phospholipase for Shp2_2SH2_ by 2.5-fold compared to the non-phosphorylated state (K_d_ ~ 0.48 ± 0.04 µM; fig. S4A and table S1). This increase in affinity is not the result of additional interactions between pY residues on Plcγ1 and Shp2_2SH2_ (fig. S4, E to H and table S1)

The existence of direct pairwise interactions between pFGFR2/Shp2 and Shp2/pPlcγ1 maintained via non-exclusive surfaces suggests the possibility of a ternary complex formation, which we postulate is the species relevant for droplet formation. Consistent with reported data (*14, 15*), we observed initial binding between pFGFR2 and Plcγ1_2SH2_ that declined as the phospholipase became phosphorylated by the receptor (Fig. 3A lanes 1-9 and fig. S5, A to C). We also observed Shp2_2SH2_ binding to pFGFR2_Cyto_ throughout the time course of the experiment (Fig. 3A lanes 10-18). When Shp2_2SH2_ and pPlcγ1_2SH2_ were mixed with FGFR2, both were found to bind the receptor concurrently (Fig. 3A lanes 19-27). Collectively, the data suggest that pFGFR2, Shp2 and pPlcγ1 form a ternary complex.

**Fig. 3.**
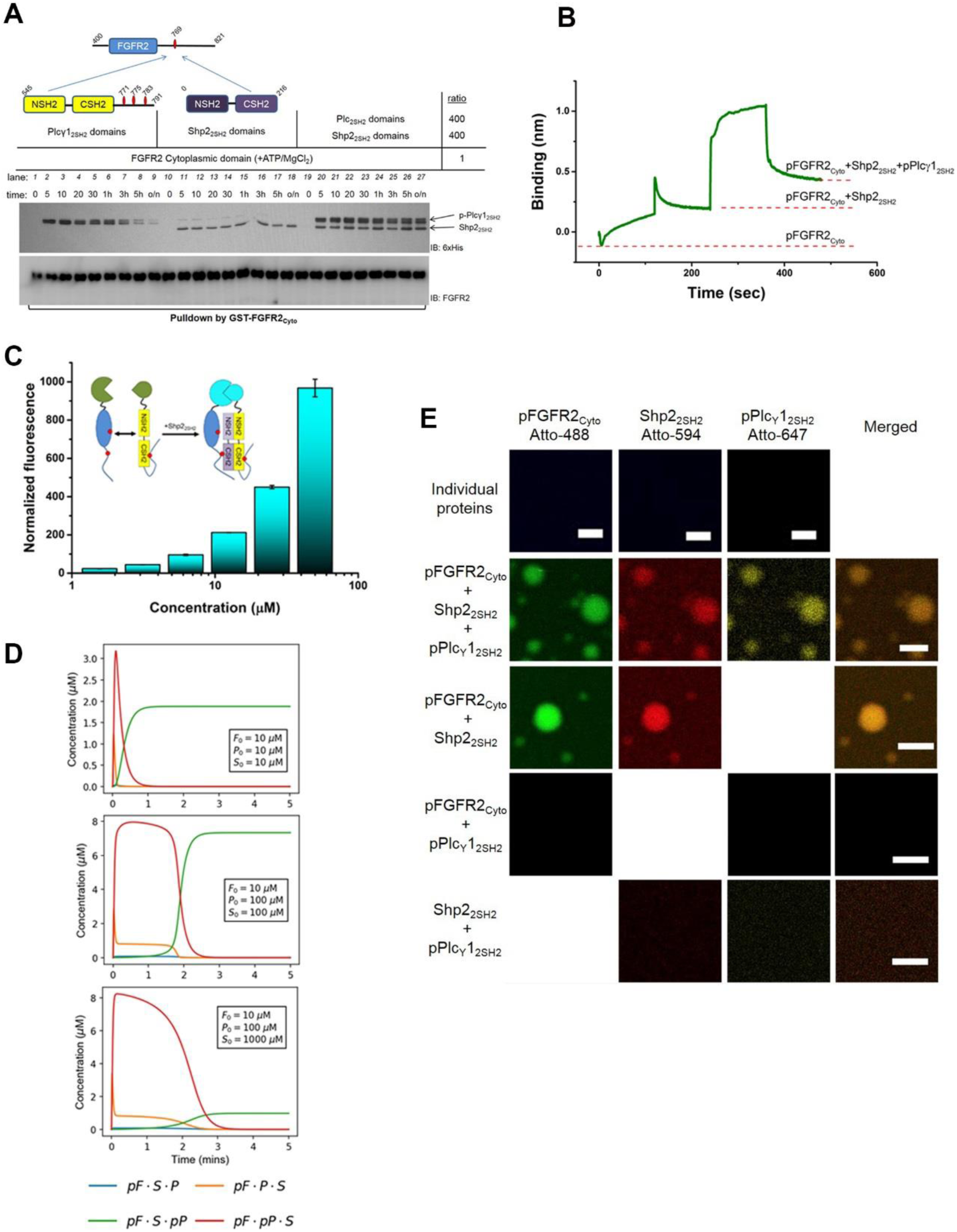
Characterisation of the ternary complex formation. (**A**) GST-pFGFR2 was used to precipitate Plcγ1_2SH2_, lanes 1-9; Shp2_2SH2_, lanes 10-18, and with both Shp2_2SH2_ and Plcγ1_2SH2_, lanes 19-27 as indicated by schematic representations of polypeptides (red ovals – pY sites). (**B**) The ternary complex was reconstituted using BLI in the presence of ATP/Mg^2+^. (**C**) BiFC was used to study the formation of ternary complex. Inset: schematic depicting the interaction: pFGFR2_Cyto_ (blue) and pPlcγ1_2SH2_ (yellow) with split CFP tag (green). Addition of increasing concentration (x-axis on graph) of Shp2_2SH2_ (purple) produces fluorescent signal (cyan). (**D**) Time evolution of the ternary complexes simulated from the deterministic mathematical model, out the four possible ternary complexes (pF·S·P, pF·P·S, pF·S·pP and pF·S·pP), only pF·S·pP prevails. (**E**) *in vitro* phase separation assay using Atto-labelled pFGFR2_Cyto_ (280 µM) and truncated Shp2_2SH2_ (700 µM) and pPlcγ1_2SH2_ (250 µM). Scale bar = 10 µm.

The mechanism of ternary complex formation was elucidated by reconstituting the interactions on a BLI sensor with immobilized GST-pFGFR2_Cyto_ (Fig. 3B). Saturating the sensor with Shp2_2SH2_ in the presence of ATP/Mg^2+^ showed binary complex formation. Subsequent exposure of GST-pFGFR2_Cyto_/Shp2_2SH2_ binary complex to excess pPlcγ1_2SH2_ which was phosphorylated by the receptor *in situ* resulted in the increase in signal indicating the recruitment of the phospholipase to the binary complex. The absence of ATP/Mg^2+^ and hence the inability of FGFR2 to release Plcγ1_2SH2_ after phosphorylation of Y783 compromised ternary complex formation (fig. S5D). Additional detail of ternary complex formation was provided using biomolecular fluorescence complementation (BiFC) where pFGFR2_Cyto_ and pPlcγ1_2SH2_ were N-terminally tagged with a split cyan fluorescent protein (CFP; CN173-pFGFR2_Cyto_ and CC173-pPlcγ1_2SH2_ respectively). We observed that pPlcγ1_2SH2_ was unable to bind to FGFR2, hence the absence of fluorescent response (Fig. 3C). However, addition of Shp2_2SH2_ resulted in fluorescent signal increase as pFGFR2_Cyto_ and pPlcγ1_2SH2_ were brought into proximity through the ternary complex formation. In addition, a high molecular weight complex which included the ternary complex was isolated using size exclusion chromatography (fig. S5E). A deterministic mathematical modelling further supports our mechanistic proposal whereby ternary complex formation is based on the pFGFR2-Shp2 complex which recruits pPlcγ1 (Fig. 3D).

Having identified the interactions which sustain the ternary complex we were able to reconstitute the features of LLPS using truncated Shp2_2SH2_ and pPlcγ1_2SH2_ (Fig. 3E and fig. S6, A and B). As with the intact proteins, pFGFR2_Cyto_, Shp2_2SH2_ and pPlcγ1_2SH2_ are sufficient for droplet formation. Reduction of the concentration of the receptor resulted in shrinking of droplet size (fig. S6C), and pre-treatment with a general phosphatase (calf intestine alkaline phosphatase, CIP) resulted in dissipation of droplets (fig. S6D). Inhibition of Shp2 binding to the receptor by incubating with a pY769-containing phosphopeptide, resulted in a reduction in the average size of droplets (fig. S6E). Again, both 1,6-hexanediol and excess of NaCl (fig. S6, F and G) were able to dissolve the pFGFR2_Cyto_/Shp2_2SH2_ droplets. FRAP also displayed the characteristics of LLPS (fig. S6H). Finally, we used the pY-binding incompetent SH2 domain mutations in Shp2_2SH2_ to show that both Shp2_2SH2_ R32A and Shp2_2SH2_ R32/138A were able to abrogate LLPS because they cannot bind to pY769 of pFGFR2_Cyto_ (fig. S6I). Importantly, Shp2_2SH2_ R138A also abrogated phase separation, suggesting that the non-specific weak interaction with the kinase domain (described in fig. S3E) is necessary for phase separation (fig. S6J).

To demonstrate a potential functional benefit of LLPS formation in the context of FGFR2 signalling, we employed *in vitro* enzymatic assays. Increased kinase activity was demonstrated through mixing an excess of the inactive K517I mutated FGFR2_Cyto_ (FGFR2_Cyto K517I_), with an LLPS containing the pFGFR2_Cyto_-Shp2_C459S_ complex. The pFGFR2_Cyto_ in the droplet efficiently phosphorylated the FGFR2_Cyto K517I_ substrate. However, dissolution of the droplet by addition of 1,6-hexanediol led to a reduction in kinase activity (Fig. 4A). The lipolytic activity of Plcγ1 was also greatly enhanced when it is in the phase-separated ternary complex compared to the isolated component proteins (Fig. 4B). Moreover, condensed droplet formation may help to buffer the effects of non-specific phosphatases on the activated receptor and pPlcγ1. (fig. S7, A and B). Finally, we demonstrated that within LLPS the activity of Shp2 toward both pFGFR2_Cyto_ and phosphopeptide substrate is lowered (Fig. 4C and fig. S7C). Thus, within the LLPS state the functional output of the ternary complex is enhanced through increased kinase and phospholipase activity, and down-regulation of the phosphatase.

**Fig. 4.**
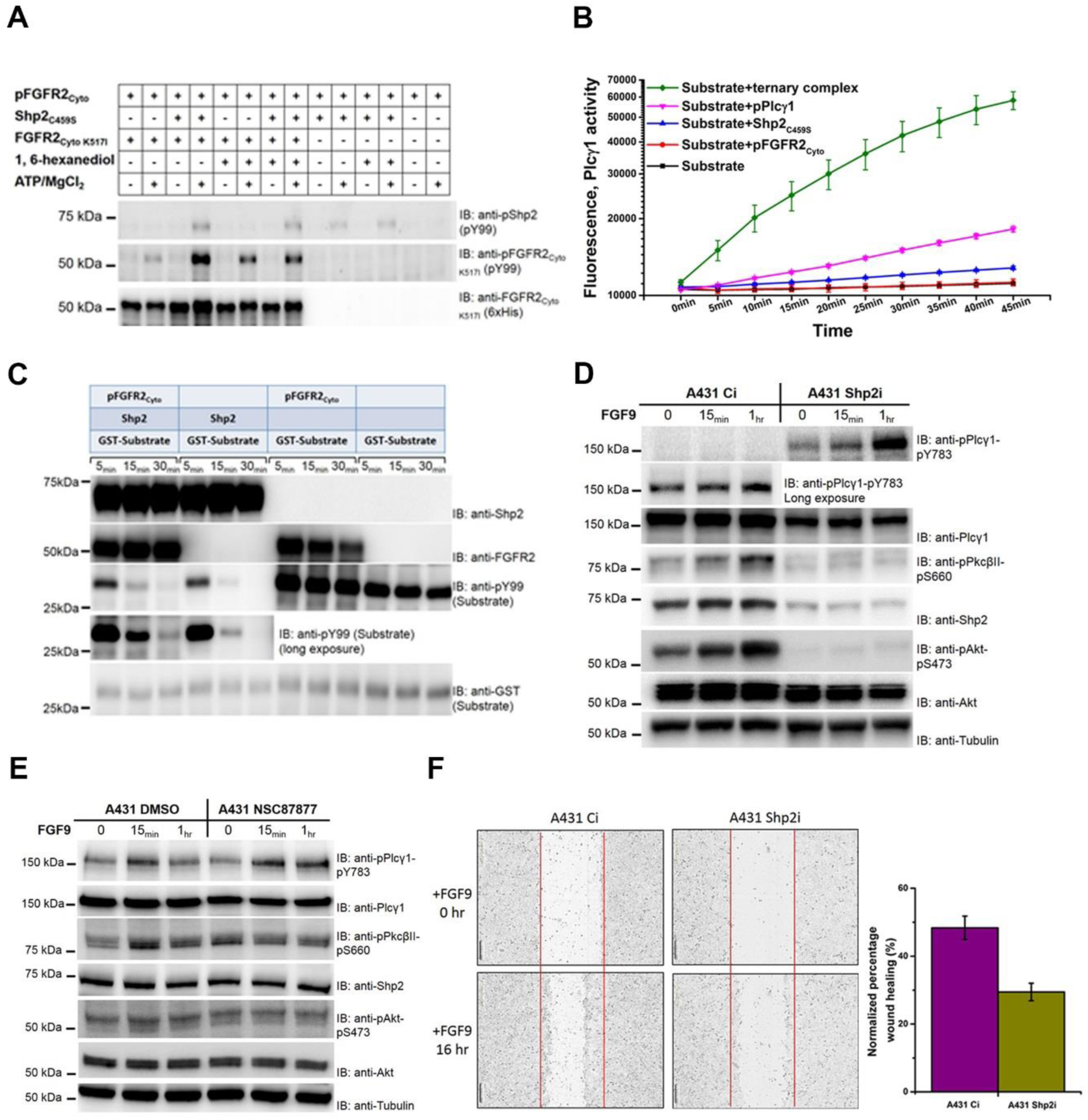
Phase transition of FGFR2-Shp2-Plcγ1 up-regulates downstream signalling. (**A**) An excess of inactive FGFR2_Cyto K517I_ was used as the substrate to monitor kinase activity. In the context of phase separated droplets, the kinase activity of pFGFR2_Cyto_ was enhanced (compare lanes 2 and 4). The addition of 1,6-hexanediol results in the reduction of kinase activity (lane 8). (**B**) Intact Plcγ1 activity was dramatically enhanced in the phase-separated environment. (n = 4) (**C**) Shp2 activity is reduced in the context of phase separated droplets compared to the isolated phosphatase (compare lanes 1-3 with 4-6 anti-pY99-identified substrate). (**D**) Depletion of Shp2 upregulates Plcγ1 through phosphorylation of Y783, but downregulates its downstream effectors (shown by reduced phosphorylation of PkcβII-S660 and Akt-S473). (**E**) Shp2 inhibitor NSC87877 has no effect on the markers for Plcγ1 signalling (phosphorylation of PkcβII-S660 and Akt-S473). (**F**) Cell motility is reduced in the absence of Shp2 in A431 cells (Shp2i) compared to control (Ci) cells. Inset shows graphical representation of percentage recovery. (n = 3)

To confirm that our *in vitro* functional studies extend to *in cellulo* context, we stably knocked down Shp2 expression in A431 cells (Shp2i). In the absence of Shp2, FGF9 stimulated cells displayed impaired dephosphorylation of Plcγ1 (Fig. 4D). Although knockdown of Shp2 should result in prolonged activation of the receptor, the observation that the phosphorylation of downstream effectors PkcβII and Akt is significantly reduced suggests that Plcγ1 function is compromised. Despite being in a phosphorylated state Plcγ1 is unable to access its membrane-localised substrate in the absence of the scaffolding function of the FGFR2-Shp2 complex. Similar results were observed using CRISPR to knock out Shp2 expression in MCF7 cells (fig. S7D). The decoupling of the phosphatase activity of Shp2 and Plcγ1-mediated signalling is further exemplified with addition of the Shp2 inhibitor NSC87877 (50 µM) to either A431 cells (Fig. 4E) or FGFR2_ΔVT_-expressing HEK293T cells (fig. S7E). In both cell lines the levels of phosphorylated PkcβII and Akt remain unaffected by inhibition of Shp2 underscoring the idea that it is the presence, rather than the activity of Shp2 which is required for Plcγ1 activity. Further corroboration of this was shown by knocking down of Shp2 in HEK293T cells that stably expressed FGFR2_ΔVT_ which led to depressed Ca^2+^ concentration (fig. S7F). In agreement with this, we also observed reduced cell motility in A431 cells (Fig. 4F) and FGFR2_ΔVT_-expressing HEK293T cells (fig. S7G), concomitant with the loss of Plcγ1 function.

In the cytoplasmic milieu proteins diffuse randomly to their receptor targets largely unaided energetically. Thus, it is expected that the propagation of a mutually exclusive signal needs to be supported by mechanisms which ensure timely delivery of the correct proteins to the RTK with greater reliability than the probability-dependent diffusion process. Collectively our data argue for a model in which the condensation of pFGFR2, Shp2 and pPlcγ1 into a LLPS state (fig. S8). The assembly of the three proteins forming the condensate are reproduced in a ternary complex suggesting that this forms the basic element of the LLPS state supported by multiple interactions between pY sites on FGFR2 and the SH2 domains of Shp2 and pPlcγ1.

A unique aspect of the LLPS complex observed in this work is that, unlike previously reported membraneless organelles or particles (*18–23*), it is sustained by a plasma membrane-bound receptor. This membrane association limits the extent of expansion of the droplet into the cytoplasm (*6, 9*), and, as a result, the extent of the LLPS state likely resembles a droplet of condensed protein on the inner leaflet of the membrane (*9, 11*). Additional work is needed to further understand biological roles of LLPS in RTK signalling, as well as investigate potential links to the manifold disease states associated with perturbations of RTK function. Here, it will be of interest to examine how LLPS formation affects efficacy of therapeutic agents that target RTKs or their effectors.

## Materials and Methods

### Cell culture

HEK293T, A431 and MCF7 cells were maintained in DMEM (Dulbecco’s modified Eagle’s high glucose medium) supplemented with 10% (v/v) FBS (foetal bovine serum) and 1% antibiotic/antimycotic (Lonza) in a humidified incubator with 10% CO_2_. shRNA control cells (A431 Ci) and Shp2 knockdown cells (A431 Shp2i) were maintained as described previously (*27*).

### Cloning, expression and purification of recombinant proteins

FGFR2 and Shp2 prokaryotic expression plasmids have been described in detail elsewhere (*27–29*). The full-length Plcγ1 expression vector was a kind gift from Dr. Matilda Katan (University College London, UK). Histidine-tagged fusion proteins were purified from BL21(DE3) cells. A single colony was used to transform 100 mL of LB which was grown overnight at 37 °C. 1L of LB were inoculated with 10 mL of this overnight culture and were allowed to grow at 37 °C until the OD_600_=0.8 at which point the culture was cooled down to 20°C and expression was induced with 1 mM IPTG. Cultures were allowed to grow for a further 12 hours before harvesting by centrifugation. Cells were re-suspended in 20 mM Tris, 150 mM NaCl, 10% glycerol, pH 8.0 in the presence of protease inhibitors and lysed by sonication. Insoluble material was removed by centrifugation (40,000 *g* at 4 °C for 60 min). The soluble fraction was applied to a Talon column. Following a wash with 10 times column volume of buffer (20 mM Tris, 150 mM NaCl, pH 8.0) protein was eluted from the column with 150 mM imidazole and was concentrated to 5 mL and applied to a Superdex75 gel filtration column in buffer containing 20 mM HEPES, 150 mM NaCl and 1 mM TCEP pH 7.5. Analysis of pure proteins on SDS-PAGE showed greater than 98% purity.

Expression of ^1^H, ^15^N, ^13^C-labelled protein for backbone resonance assignment was done as previously described (*18*). For expression in 100% D_2_O, this procedure was modified by pre-growing the culture in a small volume of 100% D_2_O prior to expression over 20 hours.

### *In vitro* phosphorylation of purified proteins

Purified proteins were phosphorylated by incubating with recombinant FGFR2_pCyto_ conjugated on agarose beads with 5 mM ATP and 10 mM MgCl_2_. FGFR2_pCyto_ protein was removed after the phosphorylation reaction by centrifugation. The phosphorylation reactions were quenched by adding EDTA (prepared in 10 mM HEPES, pH 7.5) to a final concentration of 50 mM. Proteins were analysed by SDS-PAGE and western blot to study the phosphorylation status.

### Protein fluorescent labelling

Purified proteins were labelled with Atto-488, Atto-594 and Atto-647488 (Sigma), according to the manufacturer’s protocol using a 1:1 protein:fluorescent dye mole ratio. Labelling was performed in labelling buffer (bicarbonate buffer, 0.1 M, preferably of pH 8.3). Excess label was removed G-15 desalting chromatography. Protein concentration and labelling efficiency were determined according to manufacturer’s protocol.

### In vitro droplet formation

Purified proteins were mixed according to the experimental requirement. Protein mixtures were incubated at room temperature for 1 minute unless otherwise indication before imaging.

### Confocal microscopy

Images were acquired either on Leica SP8 or Zeiss LSM880 microscopes. Samples were prepared in buffer (20mM HEPES, pH 7.5; 100 Mm NaCl, and 1mM TCEP) at indicated concentrations, and incubated at room temperature before imaging.

### Pulldown and western blots

For western blotting cells were grown in 10cm dishes, serum starved overnight and stimulated with 10ng/ml FGF9 for the indicated time period. Cells were lysed with buffer containing 50 mM Hepes (pH 7.5), 0.1% (v/v) NP-40, 10 mM NaF, 1 mM sodium orthovanadate, 10% (v/v) glycerol, 150 mM NaCl, 1 mM PMSF and Protease Inhibitor Cocktail Set III (Calbiochem). The cell debris was removed by centrifugation at 13,000 rpm for 20 minutes. The detergent soluble fraction was used for western blotting or pulldown experiments. For immunoblotting, proteins were separated by SDS-PAGE, transferred to PVDF membranes and incubated with the specific antibodies. Immune complexes were detected with horseradish peroxidase conjugated secondary antibodies and visualized by enhanced chemiluminescence reagent according to the manufacturer’s instructions (Pierce). For pulldown experiments, purified proteins or 1 mg of whole cell lysate was prepared in 1 ml volume. Proteins were immobilized on Glutathione Sepharose (GE Healthcare Life Science) or StrepTactin™ Sepharose (GE Healthcare Life Science) was added and incubated at 4 °C overnight with gentle rotation. The beads were then spun down at 4,000 rpm for 3 minutes, supernatant was removed and the beads were washed with 1 ml buffer. This washing procedure was repeated five times in order to remove non-specific binding. After the last wash, 50 µl of 2x Laemmli sample buffer were added, the sample was boiled and subjected to SDS-PAGE and western blot assay.

### Bimolecular fluorescence complementation (BiFC)

The sequences encoding resides 1-173 (CN173) and 174-238 (CC173) of CFP were fused upstream of sequences encoding FGFR2_Cyto_ and Plcγ1_2SH2_ in pET28b vectors for expression in *E. coli* (*30*). Both proteins were purified and phosphorylated as described above in the presence of FGFR2_Cyto_, ATP and MgCl_2_. The fluorescence complementation was measured using the Monolith NT.115 (NanoTemper Technologies, GmbH) as described above. Protein samples were filled into capillaries and the change in fluorescence intensity upon the adding of Shp2_2SH2_ domain was monitored. Data were plotted using Origin 7.0.

### Bio-layer interferometry (BLI)

BLI experiments were performed using a FortéBio Octet Red 384 using Anti-GST sensors. Assays were done in 384 well plates at 25 °C. Association was measured by dipping sensors into solutions of analyte protein (FGFR2 proteins) for 125 seconds and was followed by moving sensors to wash buffer for 100 seconds to monitor the dissociation process. Raw data shows a rise in signal associated with binding followed by a diminished signal after application of wash buffer. In experiments in which the ternary complex was reconstituted, GST-pFGFR2_Cyto_ was captured on a sensor, the sensor was saturated with Shp2_2SH2_ forming a binary complex as shown by the increase in signal. The binary complex was then exposed to excess Plcγ1_2SH2_. The further increase in signal on sequential addition of the two proteins to the immobilized GST-pFGFR2_Cyto_ is consistent with ternary complex formation which involves the binding of pPlcγ1_2SH2_ to the pre-existing complex of FGFR2-Shp2_2SH2_. ATP/Mg^2+^ was added in the experiment described in Fig. 3B in order to phosphorylate Plcγ1_2SH2_ by the receptor *in situ*. No ATP/Mg^2+^ was used in the experiment described in the fig. S5B.

### Microscale thermophoresis (MST)

The binding affinities were measured using the Monolith NT.115 (NanoTemper Technologies, GmbH). Proteins were fluorescently labelled with Atto488 according to the manufacturer’s protocol. Labelling efficiency was determined to be 1:1 (protein:dye) by measuring the absorbance at 280 and 488 nm. A 16 step dilution series of the unlabelled binding partner was prepared and mixed with the labelled protein at 1:1 ratio and loaded into capillaries. Measurements were performed at 25 °C in 20 mM HEPES, 150 mM NaCl and 1 mM TCEP pH 7.5 buffer containing 0.01% Tween 20. Data analyses were performed using Nanotemper Analysis software, v.1.2.101 and were plotted using Origin 7.0. All measurements were conducted as triplicates and the errors were presented as the standard error of the triplicates. Equilibrium dissociation constants (K_d_’s) are reported as ‘apparent’ values because for the interaction of FGFR2_pCyto_ with Shp2 and Plcγ1 the MST binding profiles appear to include two independent binding sites. The second of which is much weaker and is likely to be due to non-specific effects. Nonetheless the possibility of competing equilibria is flagged by the use of the term apparent. Reported data fits are based only on the initial tight binding event. Where the K_d_ is reported without the superscript ‘app’ the data has been fit to the standard 1:1 binding model.

### Isothermal titration calorimetry (ITC)

ITC experiments were carried out using a MicroCal iTC200 (Malvern) at 25 °C. 20 15 µl injections of 100 µM FGFR2_pC58_ were made into 10 µM Shp2_2SH2_ in the calorimeter cell. A control experiment involving the injection of 100 µM FGFR2_pC58_ into buffer was performed. The heat per injection was determined and subtracted from the binding data. Data was analysed using a single independent site model using Origin software.

### Nuclear magnetic resonance (NMR)

All NMR spectroscopic experiments were carried out on Bruker Avance III (700, 800 and 950 MHz) NMR spectrometers equipped with cryogenically cooled triple resonance probes with a z-axis pulse field gradient coil were used. Resonance assignment spectra were recorded in 25 mM Na_2_HPO_4_/NaH_2_PO_4_, pH = 6.5, 50 mM NaCl, 5 mM DTT, 1 mM EDTA. For the ^1^H,^15^N,^13^C labelled construct, a standard set of 3D backbone resonance assignment experiments (HNCA, HNCOCA, HNCACB, CACBCONH, HNCO and HNCACO) using standard Bruker library pulse sequences (with Watergate water suppression) or BEST versions of amide transverse relaxation optimized spectroscopy (TROSY) (*31*) pulse sequences applying Non-Uniform Sampling (17-25%) were used to obtain a high-resolution spectra.

The NMR titration of Shp2 into Plcγ1 experiments were recorded at 25 ºC using 200 µM uniformly ^15^N-labelled Plcγ1_2SH2_ in 20 mM HEPES (pH 7.5) containing 150 mM NaCl, 1mM TCEP and 10% (v/v) D_2_O. 0, 150, 300, 450 600 and 900 µM unlabelled Shp2_2SH2_ were added and an amide BEST TROSY pulse sequence recorded.

All NMR data was processed with NMRPipe (*32*) and analyzed with CcpNmr Analysis software package (*33, 34*) available in-house and on NMRBox platform (*35*). Chemical shift perturbations (CSPs) for individual residues were calculated from the chemical shift for the backbone amide ^1^H (Δω_H_) and ^15^N (Δω_N_) using the following equation: 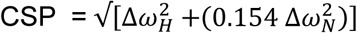 (36).

### Calcium concentration assay

The calcium assay was perform using Fluo-4 NW Calcium Assay Kit (F36206, ThermoFisher Scientific) according to the manufactural manual. Briefly, HEK293T-FGFR2_ΔVT_ cells were transiently transfected with control (scrambled) and Shp2 shRNA. Knockdown cells were seeded in a 96 well plate. After FGF9 stimulation, medium was removed and cells were exposed to the dye loading solution and incubated for 30 minutes at 37°C then 30 minutes at 25°C. The Ca^2+^ level (fluorescence) was measured using instrument settings for excitation at 490 nm and emission at 510 nm.

### In vitro Plcγ1 activity assay

The lipolytic activity of Plcγ1 was determined using the artificial substrate 4-methylumbelliferyl myo-inositol-1-phosphate, N-methyl-morpholine salt (Biosynth International Inc., Itasca, IL). Briefly, reaction mixtures consisted of 10 mM Tris, pH 6.8, 0.8 mM substrate, and 50% of protein mixtures. The reaction mixtures were placed in a 96-well plate, and fluorescence was measured using 350 nm excitation/450-nm emission filters on a plate reader. Reactions were allowed to proceed for 45 min at room temperature, with measurements taken every 5 min.

### Wound-healing assay

A431Ci and Shp2i cells were seeded in a 96 well plate. After overnight serum starvation the cells were scratched using IncuCyte® WoundMaker and stimulated with 10 ng/ml FGF9, then incubated for further 6 hr. Images were taken at 0 hr and after 6 hr by a microscope gantry inside a cell incubator (Incucyte, Essen Bioscience, Ann Arbor, MI, USA). The identical experiment was also performed using HEK293T cells stably transfected with FGFR2_ΔVT_.

## Acknowledgments

We thank Anastasia Zhuravleva, (University of Leeds, School of Molecular and Cellular Biology) for critical discussion and useful comments. We thank Dr Arnout Kalverda (University of Leeds, bioNMR Facility) for assistance with NMR data collection. Additional editing of this work was provided by Life Science Editors.

## Author contributions

C.-C.L., K.M.S., L.W., A.S., C.S., H.K., and Z.A. performed experiments and analysed data. P.-A.J performed mathematical and numerical model analysis. C.-C.L., K.M.S., E.M., and J.E.L conceived experiments, and C.M.-P. and J.E.L. developed the mathematical model. C.-C.L., K.M.S. and J.E.L. wrote the manuscript.

