## Supplemental Data for "Receptor tyrosine kinases regulate signal transduction through a liquid–liquid phase separated state"

### SUPPLEMENTARY MATERIALS

Fig. S1 to S8

Table S1

Mathematical model

Reaction Figure

### Supplementary Figure 1

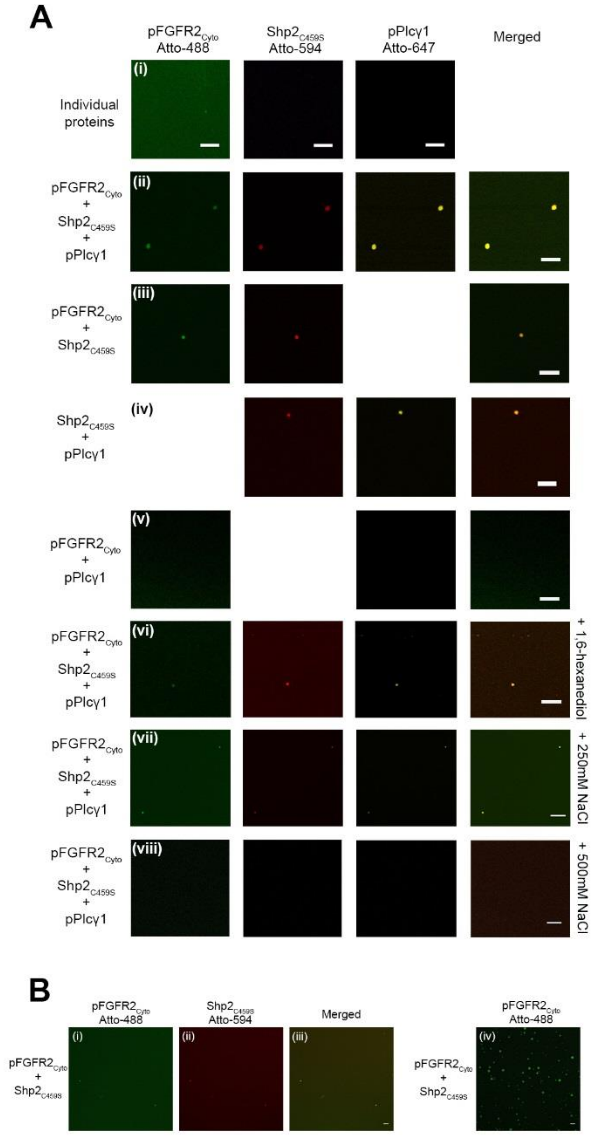


**Fig. S1. FGFR2-Shp2-Plcγ1 complex results in the formation of liquid droplets.** (**A**) Droplet formation observed between pFGFR2$_{Cyto}$ Atto-488 (6.6 µM), Shp2$_{C459S}$ Atto-594 (60 µM) and pPlcγ1 Atto-647 (15 µM). (i) Individual proteins showed no evidence of phase transition. Droplet formation was observed after 1 minute between different combinations of proteins: (ii) all three proteins; (iii) pFGFR2$_{Cyto}$ with Shp2$_{C459S}$ and (iv) Shp2$_{C459S}$ with pPlcγ1. (v) No droplet formation was observed with pFGFR2$_{Cyto}$ with pPlcγ1. Droplet size was diminished in the presence of 1, 6-hexanediol for 30 minutes (vi) and with increasing concentration of NaCl after incubation for 10 minutes (vii, viii) compared to (ii). Scale bar = 10 µm. (**B**) The presence of the fluorescent label (Atto594) on Shp2 impinges on the size of droplet. Droplets formed with $pFGFR2_{Cyto}$-Atto488 and $Shp2_{C459S}$-Atto594 after 1 minute (panels i to iii) are smaller than droplets formed in the absence of Atto594 over the identical time period (iv). Scale bar = 10 µm.

### Supplementary Figure 2

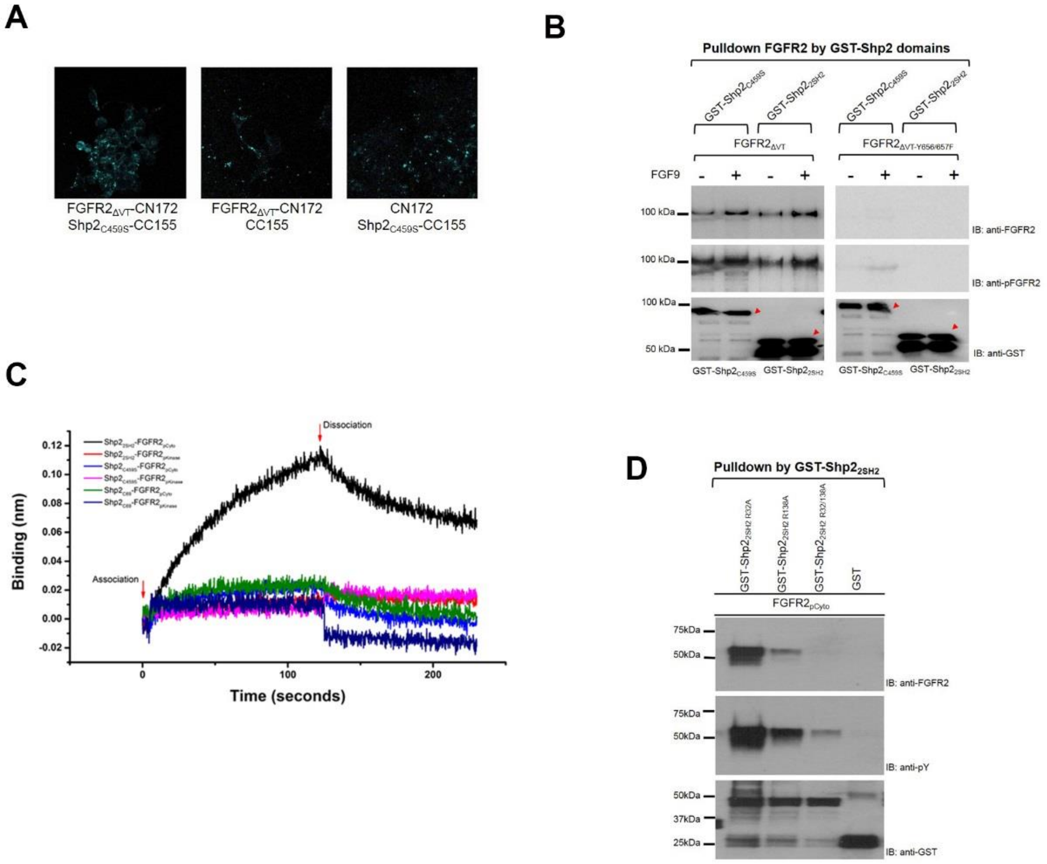


**Fig. S2. FGFR2-Shp2 interaction is tyrosine-phosphorylation-dependent.** (**A**) Reconstitution of the CFP fluorophore (from CN172 and CC155 polypeptide fusion tags on $FGFR2_{\Delta VT}$ and $Shp2_{C459S}$ respectively) in BiFC shows direct interaction between the receptor and the phosphatase in HEK293T cells. (**B**) Pull down experiments using GST-$Shp2_{C459S}$ or GST-$Shp2_{2SH2}$ show that binding of Shp2 requires phosphorylation of $FGFR2_{\Delta VT}$. $FGFR2_{\Delta VT}$ or $FGFR2_{\Delta VT\text{-}Y656/657F}$ were transfected into HEK293T cells which were unstimulated or FGF9-stimulated. The red arrows highlight the GST fusion as part of the Shp2 constructs. (**C**) BLI experiments were used to confirm the phosphorylation-dependent interaction and the specific domains required for binding. GST-$Shp2_{2SH2}$, GST-$Shp2_{C459S}$ and GST-$Shp2_{C69}$ were immobilized on GST sensors and $pFGFR2_{Cyto}$ and $pFGFR2_{kinase}$ were used to test the binding.

$pFGFR2_{Cyto}$ clearly interacts with GST-$Shp2_{2SH2}$. The expected weak binding to $pFGFR2_{kinase}$ is not visible due to the concentration of this reagent being below the $K_d$ for the interation. Black - immobilized $Shp2_{2SH2}$ with $pFGFR2_{Cyto}$; red - immobilized $Shp2_{2SH2}$ with $pFGFR2_{kinase}$; blue - immobilized $Shp2_{PTPC459S}$ with $pFGFR2_{Cyto}$; pink - immobilized $Shp2_{PTPC459S}$ with $pFGFR2_{kinase}$; green - immobilized $Shp2_{C69}$ with $pFGFR2_{Cyto}$; dark blue - immobilized $Shp2_{C69}$ with $pFGFR2_{kinase}$. Ligand-analyte association (0 sec) and dissociation (buffer washing, 120 sec) are indicated by red arrows. (**D**) GST-$Shp2_{2SH2}$, GST-$Shp2_{2SH2\ R32A}$, GST-$Shp2_{2SH2\ R138A}$ and GST-$Shp2_{2SH2\ R32/138A}$ were used to pull down $pFGFR2_{Cyto}$. Mutation of R138 abrogates binding of FGFR2 confirming the requirement of a wild type CSH2 domain for binding to receptor.

### Supplementary Figure 3

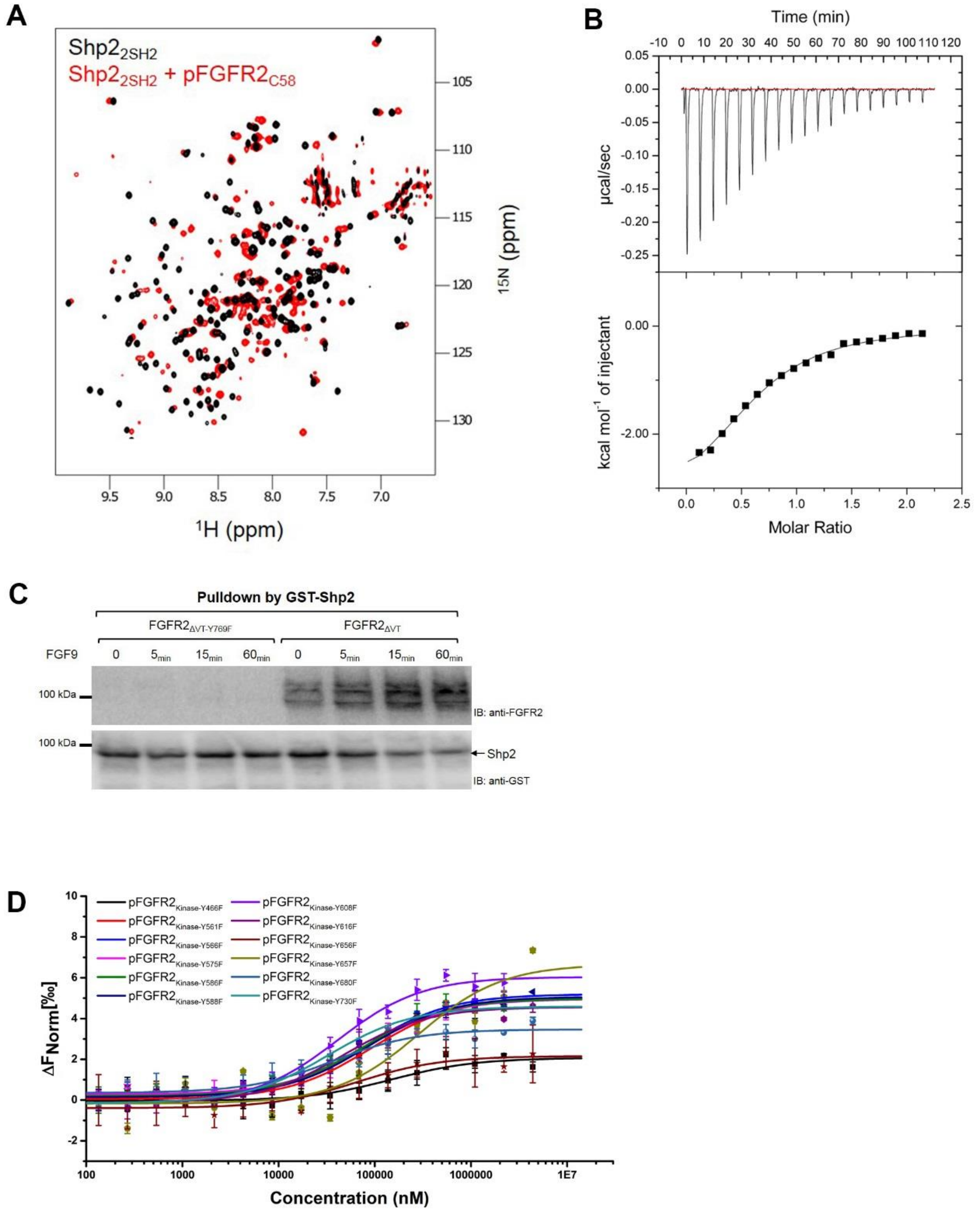


**Fig. S3. Different binding of the phosphorylated FGFR2 C-terminal tail and the kinase domain to Shp2.** (**A**) NMR spectra of $^{1}$H, $^{15}$N-labelled isolated Shp2$_{2SH2}$ (black) and with added pFGFR2$_{C58}$ (red). The chemical shifts confirm direct interaction across a broad interface. (**B**) Binding of Shp2$_{2SH2}$ with the pFGFR2$_{C58}$ shown by ITC. The top panel shows raw data for the titration; the bottom panel shows integrated peaks plotted on axes with molar heat of binding versus the molar ratio of titrated protein fitted to a single-site binding model.

Heats of dilution were measured in a separate control experiment and subtracted from binding data prior to fitting. Importantly the stoichiometry of the interaction is 1:1 confirming that only CSH2 from the tandem SH2 domains recognizes pY769 on the receptor. (**C**) Pull down experiment using GST-Shp2$_{C459S}$. FGFR2$_{\Delta VT}$ and FGFR2$_{\Delta VT\text{-}Y769F}$ were transfected into HEK293T cells which were unstimulated or FGF9-stimulated. GST-Shp2$_{C459S}$ binds to FGFR2$_{\Delta VT}$, but not to FGFR2$_{\Delta VT\text{-}Y769F}$ confirming the interaction is mediated by pY769. (**D**) MST measurements of Shp2$_{2SH2\ R138A}$ binding to pFGFR2$_{kinase}$ with single Y to F mutants of all of the individual tyrosines on the kinase domain. This result shows multivalent, weak binding between Shp2 NSH2 domain and any of the available pY residues.

### Supplementary Figure 4

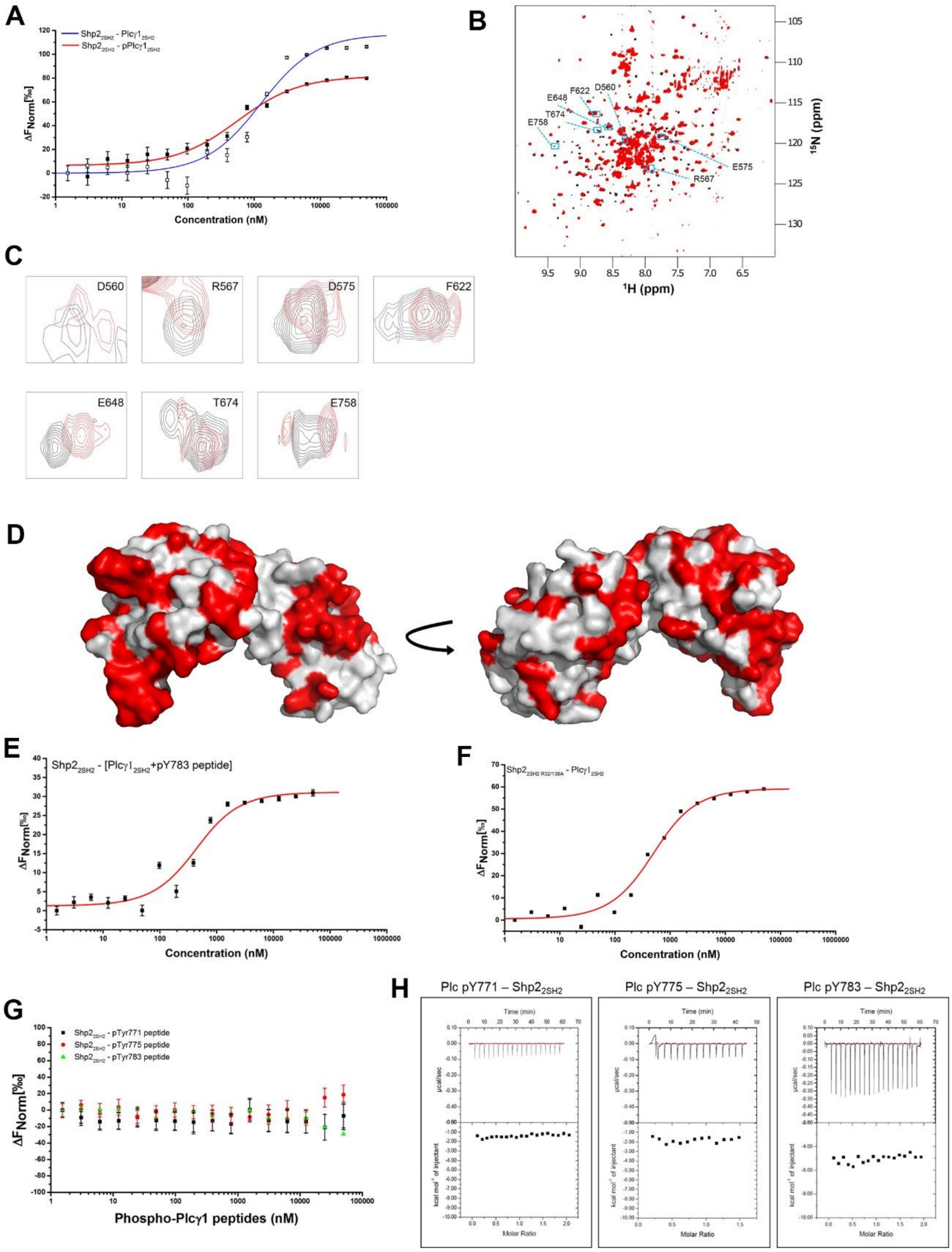


**Fig. S4. Phosphorylation-independent Shp2$_{2SH2}$ and Plcγ1$_{2SH2}$ interaction.** (**A**) MST measurement of Shp2$_{2SH2}$ binding to labelled Plcγ1$_{2SH2}$, blue curve; or labelled pPlcγ1$_{2SH2}$, red curve. (**B**) NMR $^{1}$H and $^{15}$N chemical shift changes on addition of Shp2$_{2SH2}$ to $^{15}$N-labelled

Plcγ1$_{2SH2}$. Blue squares highlight some of the residues on the spectrum showing shift changes; which are magnified in (**C**). (**D**) $^{1}$H, $^{15}$N peak assignments mapped onto the space-filling model crystal structure of Plcγ1$_{2SH2}$ (PDB code: 4FBN). The orientations of the structure are as shown in Fig. 2e. The coverage of assigned residues of Plcγ1$_{2SH2}$ is 46.5% (red, assigned residues). (**E**) MST isotherm for the interaction of labelled Shp2$_{2SH2}$ and a preformed complex between Plcγ1$_{2SH2}$ and a tyrosyl phosphopeptide containing pY783 showing that the binding of pY783 does not hinder the tandem SH2 domain interface. (**F**) MST isotherm for the interaction of labelled Shp2$_{2SH2\ R32/138A}$ and pPlcγ1$_{2SH2}$, showing that the interaction is not based on the canonical binding of pY to an SH2 domain. (**G**) MST isotherm for the binding of synthesized Plcγ1-derived tyrosyl phosphopeptides containing pY771, pY775 or pY783 to labelled Shp2$_{2SH2}$. No significant interaction was found with any of the phosphopeptides. (**H**) ITC measurements of Shp2$_{2SH2}$ binding to Plcγ1 pY771, pY775, and pY783 tyrosyl phosphopeptides. Twenty 3 µl injections of each phosphopeptide (100 µM) were titrated into Shp2$_{2SH2}$ (10 µM) at 25°C. Top, baseline-corrected power-versus-time plot for the titration. Bottom, integrated heats and curve fitting using Origin™ software.

### Supplementary Figure 5

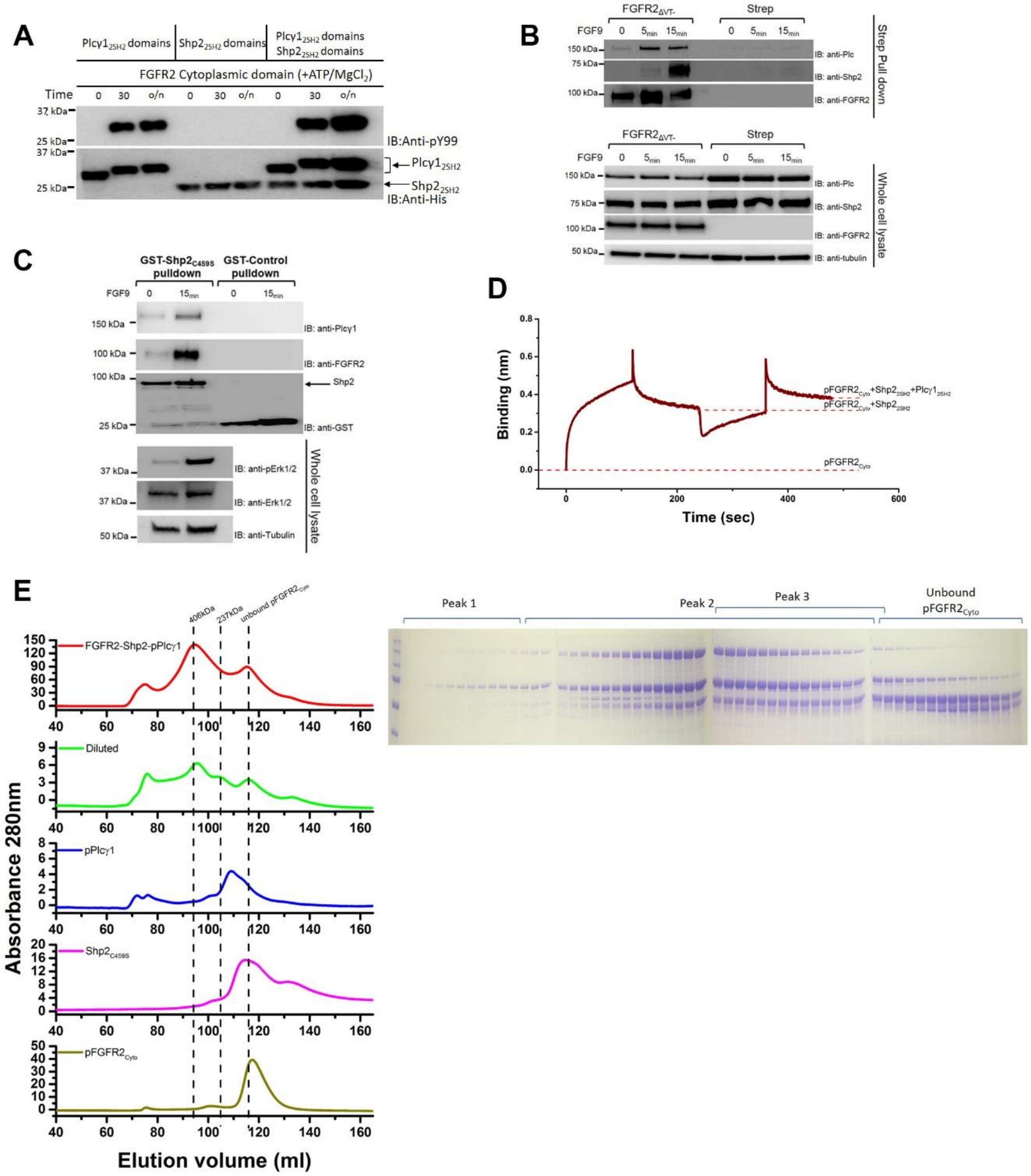

**Fig. S5. The formation of $pFGFR2_{Cyto}$-$Shp2_{C459S}$-pPlcγ1 ternary complex.** (**A**) Blot showing the efficiency of phosphorylation of $Plcγ1_{2SH2}$ and $Shp2_{2SH2}$ by $FGFR2_{Cyto}$ for proteins used in binding experiments. Recombinant proteins were mixed and incubated with ATP/$Mg^{2+}$ for specified period of time. A general anti-phosphotyrosine antibody (pY99) was used to examine the phosphorylation level, and an anti-His-tag antibody was used to confirm the total protein level. No protein degradation was observed. (**B**) Endogenous Plcγ1 and Shp2 were both precipitated by Strep-FGFR2$_{\Delta VT}$ in HEK293T stimulated with FGF9. Relative expression levels shown in the whole cell lysate blot. (**C**) Endogenous Plcγ1 and over-expressed FGFR2$_{\Delta VT}$ were pulled down by GST-Shp2$_{C459S}$ mutant from HEK293T lysate. Whole cell lysates were probed with an anti-phospho-Erk1/2 antibody to confirm the basal and activated states. (**D**) The ternary complex cannot be reconstituted in the absence ATP/$Mg^{2+}$. Sequential binding of Shp2$_{2SH2}$ (0 sec) and Plcγ1$_{2SH2}$ (240 sec) to GST-pFGFR2$_{Cyto}$ captured on an anti-GST sensor using BLI. (**E**) Size exclusion chromatography was used to isolate the pFGFR2Cyto-Shp2C459S-pPlcγ1 complex. The components of each elution fraction were examined by coomassie gel staining. A complex of greater than 1:1:1 of pFGFR2Cyto-Shp2C459S-pPlcγ1 ternary complex was observed with a calculated molecular weight of 406 kDa (peak 2). The binary pFGFR2Cyto-Shp2C459S complex was identified as peak 3.

### Supplementary Figure 6

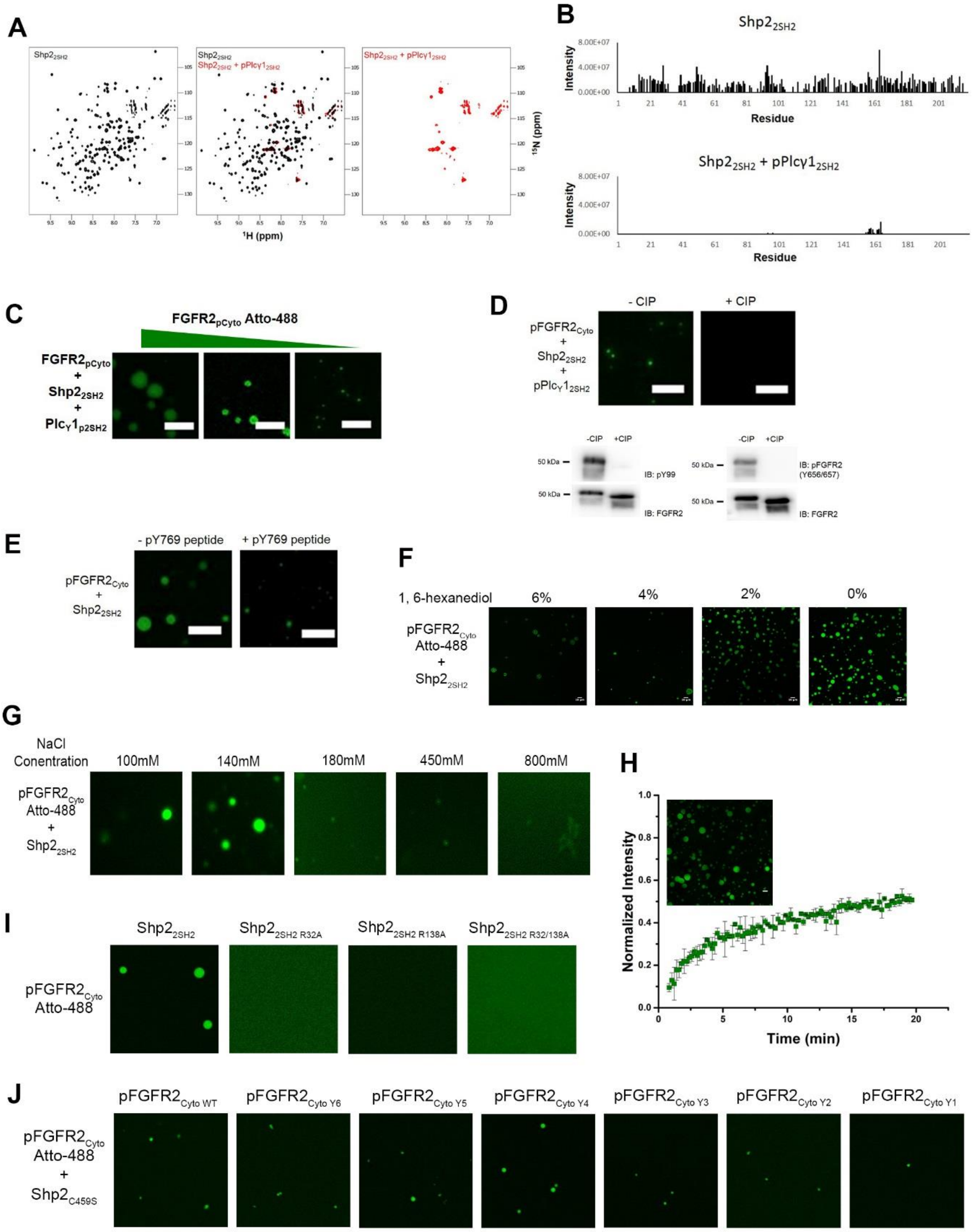


**Fig. S6. The tandem SH2 domains Shp2 and pPlcγ1 for droplets in the presence of pFGFR2$_{Cyto}$.** (**A**) and (**B**) Titration of pPlcγ1$_{2SH2}$ into $^{15}$N-labelled Shp2$_{2SH2}$ results in the disappearance of the majority of the peaks on the plot at elevated concentrations. The previously observed phase separation of pPlcγ$1_{2SH2}$ and $Shp2_{2SH2}$ results in line broadening making the peaks invisible to NMR. (**C**) Increase of $pFGFR2_{Cyto}$ concentration (0.3 µM, 1 µM, and 4 µM) enhances the size of LLPS. $pFGFR2_{Cyto}$, $Shp2_{C459S}$ (100 µM) and pPlcγ1 (100 µM) were added for 1 minute then imaged. Scale bar = 10 µm. (**D**) The requirement for phosphorylation of the receptor was shown since on incubation of $pFGFR2_{Cyto}$ (1 µM) with a general phosphatase (CIP) before adding $Shp2_{2SH2}$-pPlcγ$1_{2SH2}$ droplet formation was abrogated. Scale bar = 10 µm. (**E**) Incubation with pY769 peptide inhibits the binding of $Shp2_{2SH2}$ to $pFGFR2_{Cyto,}$ resulting in the loss of the LLPS state. Scale bar = 10 µm. (**F**) Addition of 1,6-hexanediol to phase separated $pFGFR2_{Cyto}$-Atto488 (1 µM) and $Shp2_{2SH2}$ (100 µM) dissolves droplets. (**G**) Addition of NaCl to phase separated $pFGFR2_{Cyto}$-Atto488 (1 µM) and $Shp2_{2SH2}$ (100 µM) dissolves droplets. Scale bar = 10 µm. (**H**) FRAP recovery curve for $pFGFR2_{Cyto}$-Atto488 (7 µM) and $Shp2_{2SH2}$ (100 µM). Error bars indicate standard error of the mean (n = 3). Inset shows the photobleached droplet population. (**I**) R to A mutation of residues 32 and 138 in the pY binding sites show that both wild type SH2 domains of Shp2 are required for LLPS with $pFGFR2_{Cyto}$. Scale bar = 10 µm. (**J**) Sequential Y to F mutation of the six exposed tyrosines on the kinase domain of FGFR2 shows that these droplets prevail as the multi-valency of the receptor is reduced. Scale bar = 10 µm.

### Supplementary Figure 7

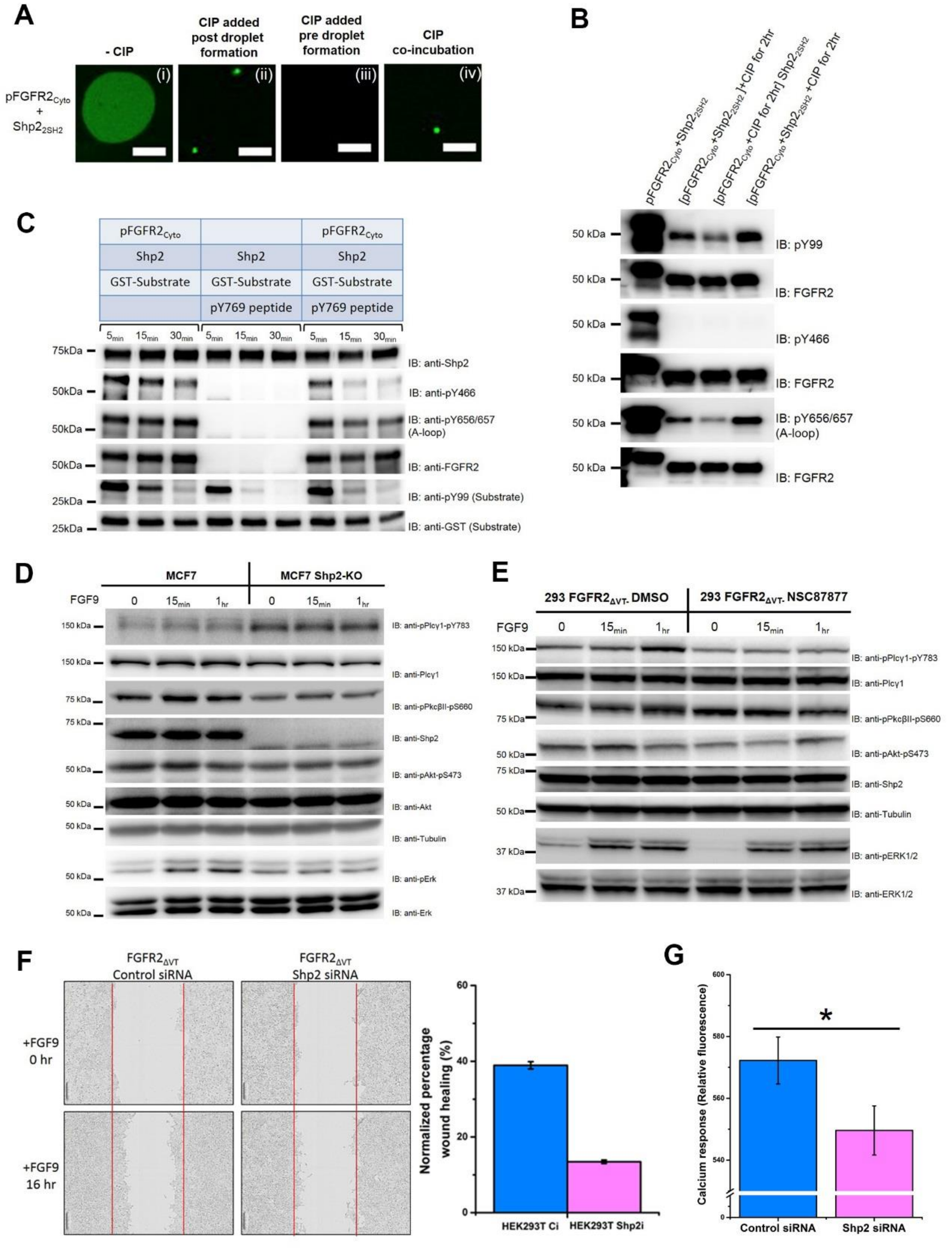


**Fig. S7. Regulation of enzymatic activities of FGFR2, Shp2, and Plcγ1 in the droplet environment.** (**A**) pFGFR2$_{Cyto}$ (50 µM)-Shp2$_{2SH2}$ (60 µM) droplet formation after 30 min. (i) is impaired on addition of general phosphatase CIP (ii), and it is completely abrogated if CIP is added initially (iii). However, co-incubation leads to rescue of the LLPS state (iv). This is consisten with the LLPS resisting dephosphorylation by phosphatase. Scale bar = 10 μm. (**B**) Western blots confirmed that pFGFR2$_{Cyto}$ (50 μM)-Shp2$_{2SH2}$ (60 μM) droplet formation after 1 min. prevents the dephosphorylation of pFGFR2$_{Cyto}$ by CIP (exposure 2 hr.), including activation loop residues pY656/657. (**C**) Using a pY769 peptide to disrupt the LLPS formation results in the restoration of Shp2 activity toward the substrate peptide (compare lanes 1-3 with 7-9). (**D**) Western blot showing MCF7 cells with knock-out of Shp2 by CRISPR (Shp2-KO). Phosphorylation of Plcγ1 Y783 is increased on FGF9 stimulation of FGFR2. Phosphorylation of S660 on PkcβII and S473 on Akt act as markers for up-regulation of Plcγ1 signalling. This is suppressed in the absence of Shp2 in the Shp2-KO cells. (**E**) Western blot showing the presence of phosphorylated downstream effector proteins in HEK293T cells without (using only DMSO vehicle) or with NSC87877 Shp2 inhibitor. The negligible change of phosphorylation of Pkc and Akt shows that Shp2 phosphatase activity does not affect Plcγ1 activity. (**F**) Wound-healing assay in HEK293T cells stably transfected with FGFR2$_{\Delta VT}$ with and without shRNA Shp2 knock down. Images from the beginning of the experiment (0 hr) and at 16 hr. Inset shows graphical representation of percentage wound healing for both cell lines; HEK293T with scrambled shRNA (light blue) and Shp2 shRNA (light pink) cells. (n = 3). (**G**) Inhibition of calcium response in Shp2 shRNA knockdown HEK293T cells (Shp2i) transfected with FGFR2$_{\Delta VT}$, compared to control cells with scrambled shRNA (Ci). *=p-value<0.002 student's t-test (n = 18).

### Supplementary Figure 8

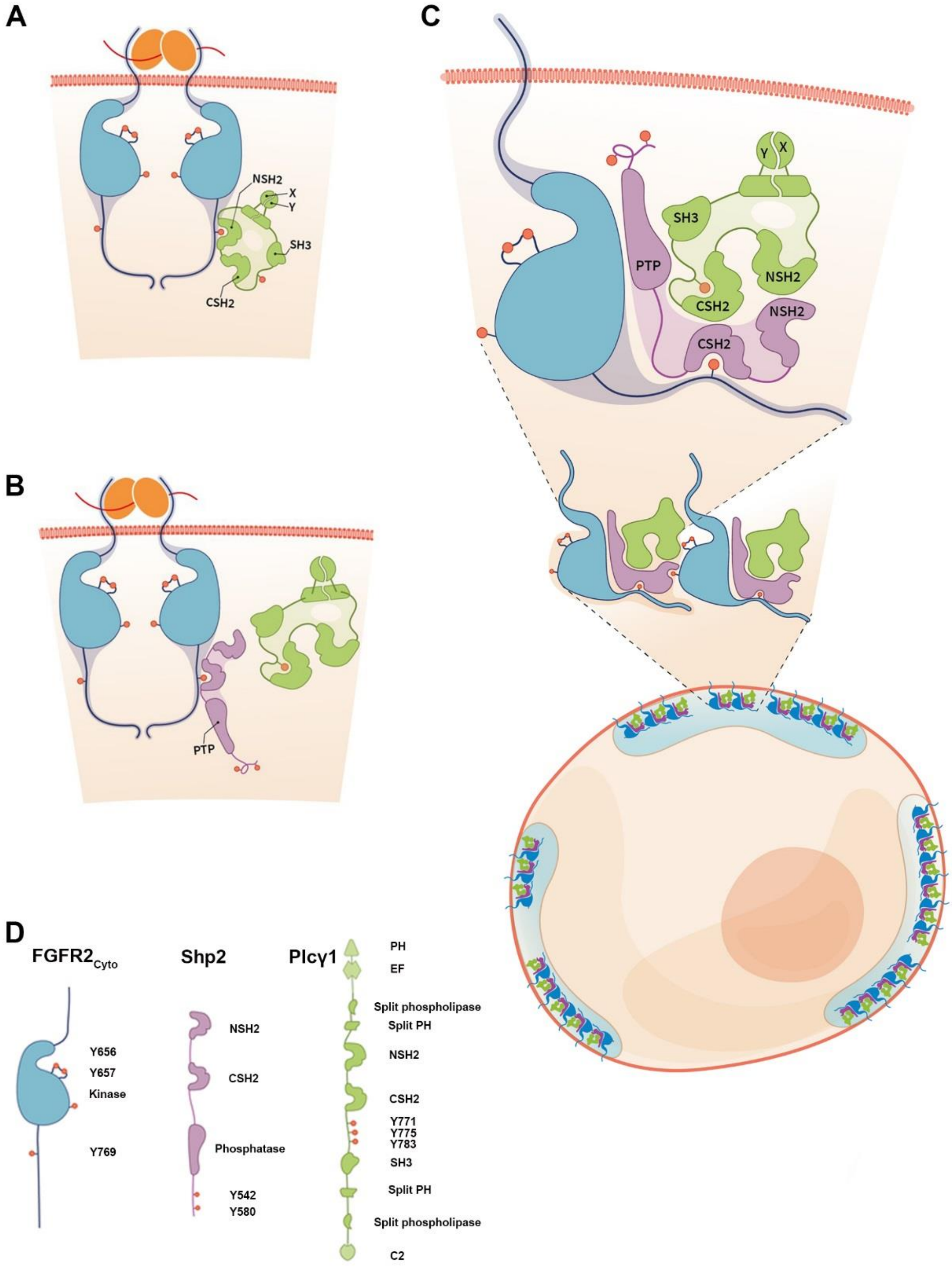


**Fig. S8. Schematic representation of formation of ternary complex in the LLPS.** (**A**) Binding of FGF/heparin (orange with red line) leads to the autophosphorylation of plasma membrane-bound receptor (blue) on numerous sites (pY – red dots). Plcγ1 (green) is recruited through its NSH2 domain to the pY769 site on the receptor tail resulting on phosphorylation of

Plcγ1 on Y783. (**B**) intermediate state whereby phosphorylation of Y783 on Plcγ1 results in an intramolecular interaction of pY783 and the CSH2 domain; this leads to a conformational rearrangement which disables pPlcγ1 from binding to the receptor. In the absence of Plcγ1 binding the pY769 site on pFGFR2 is available for binding to Shp2 (purple) through its CSH2 domain. (**C**) The pFGFR2-Shp2 complex is able to recruit pPlcγ1 via interaction between the tandem SH2 domains of Shp2 and pPlcγ1 producing the ternary complex. NSH2 on Shp2 is not occluded by the tandem SH2 domain interface. The multiple phosphorylation sites on $pFGFR2_{kinase}$ (not shown) provide secondary binding sites for Shp2 NSH2. The multivalent interactions between pFGFR2, Shp2 and pPlcγ1 allow the ternary complex to self-associated into phase separated droplets on the cellular membrane. (**D**) Schematic diagrams of the domains of $FGFR2_{Cyto}$, Shp2, and Plcγ1.

**Table S1. The dissociation constant ($K_d$) measured by MST in this study**

| **The dissociation constant ($K_d$) measured by MST in this study** | |
|---|---|
| **Protein-protein interaction** | **$K_d$ (µM)** |
| $Shp2_{2SH2\ R138A}$ – $pFGFR2_{kinase\ Y466F}$ | 151 ± 34 µM |
| $Shp2_{2SH2\ R138A}$ – $pFGFR2_{kinase\ Y561F}$ | 90.3 ± 6.1 µM |
| $Shp2_{2SH2\ R138A}$ – $pFGFR2_{kinase\ Y566F}$ | 82.6 ± 4.7 µM |
| $Shp2_{2SH2\ R138A}$ – $pFGFR2_{kinase\ Y575F}$ | 81.6 ± 5.0 µM |
| $Shp2_{2SH2\ R138A}$ – $pFGFR2_{kinase\ Y586F}$ | 74.7 ± 7.0 µM |
| $Shp2_{2SH2\ R138A}$ – $pFGFR2_{kinase\ Y588F}$ | 83.1 ± 7.1 µM |
| $Shp2_{2SH2\ R138A}$ – $pFGFR2_{kinase\ Y608F}$ | 41.3 ± 2.6 µM |
| $Shp2_{2SH2\ R138A}$ – $pFGFR2_{kinase\ Y616F}$ | 47.7 ± 3.9 µM |
| $Shp2_{2SH2\ R138A}$ – $pFGFR2_{kinase\ Y656F}$ | 65.3 ± 10.7 µM |
| $Shp2_{2SH2\ R138A}$ – $pFGFR2_{kinase\ Y657F}$ | 288 ± 45 µM |
| $Shp2_{2SH2\ R138A}$ – $pFGFR2_{kinase\ Y680F}$ | 46.4 ± 4.2 µM |
| $Shp2_{2SH2\ R138A}$ – $pFGFR2_{kinase\ Y733F}$ | 43.4 ± 2.92 µM |
| $Shp2_{2SH2}$ – $Plc\gamma1_{2SH2}$ | 1.16 ± 0.09 µM |
| $Shp2_{2SH2}$ –$pPlc\gamma1_{2SH2}$ | 0.48 ± 0.04 µM |
| $Shp2_{2SH2}$ – [$Plc\gamma1_{2SH2}$+ pY783 peptide] | 0.39 ± 0.04 µM |
| $Shp2_{2SH2,R32/138A}$ –$Plc\gamma1_{p2SH2}$ | 0.48 ± 0.03 µM |
| $Shp2_{2SH2}$ – pY771 peptide | No binding |
| $Shp2_{2SH2}$ – pY775 peptide | No binding |
| $Shp2_{2SH2}$ – pY783 peptide | No binding |

**Mathematical model.** We have developed a deterministic mathematical model to describe the *in vitro* interactions between FGFR2$_{Cyto}$ (*F*), Plcγ1$_{2SH2}$ (*P*) and Shp2$_{2SH2}$ (*S*). The model has been formulated to include the reactions depicted in Reaction Figure, which involve the following chemical species

- $F$ (or $n_1$) = unphosphorylated FGFR2,
- $pF$ (or $n_2$) = phosphorylated FGFR2,
- $S$ (or $n_3$) = Shp2,
- $pF\ \cdot S$ (or $n_4$) = phosphorylated FGFR2-Shp2 complex,
- $P$ (or $n_5$) = unphosphorylated Plcγ,
- $pF\ \cdot P$ (or $n_6$) = phosphorylated FGFR2-Plcγ complex,
- $pF\ \cdot pP$ (or $n_7$) = phosphorylated FGFR2-phosphorylated Plcγ complex,
- $pP$ (or $n_8$) = phosphorylated Plcγ,
- $S\ \cdot P$ (or $n_9$) = Shp2-unphosphorylated Plcγ complex,
- $S\ \cdot pP$ (or $n_{10}$) = Shp2-phosphorylated Plcγ complex,
- $pF\ \cdot S\ \cdot P$ (or $n_{11}$) = phosphorylated FGFR2-Shp2-unphosphorylated Plcγ complex,
- $pF\ \cdot P\ \cdot S$ (or $n_{12}$) = phosphorylated FGFR2-unphosphorylated Plcγ-Shp2 complex,
- $pF\ \cdot S\ \cdot pP$ (or $n_{13}$) = phosphorylated FGFR2-Shp2-phosphorylated Plcγ complex, and
- $pF\ \cdot pP\ \cdot S$ (or $n_{14}$) = phosphorylated FGFR2-phosphorylated Plcγ-Shp2 complex,

where in every row above the first symbol is the abbreviated species name provided in Reaction Figure and the symbol in parentheses is used in the differential equations describing the variables of the mathematical model. We have assumed that there are no allosteric binding effects between Shp2, Plcγ, or the phosphorylated receptor. From the reactions in Reaction Figure and assuming mass-action kinetics, a set of ordinary differential equations (ODEs) for the concentrations (in units of $\mu M$) for each molecular species can be written as follows:

$$\frac{dn_1}{dt} = -k_{+1}n_1, \quad (1)$$

$$\frac{dn_2}{dt} = k_{+1}n_1 - k_{+2}n_2(n_3 + n_9 + n_{10}) + k_{-2}(n_4 + n_{11} + n_{13}) - k_{+3}n_2(n_5 + n_9) +$$

$$k_{-3}(n_6 + n_{12}) + k_{+5}n_7 + k_{+5}n_{14}, \quad (2)$$

$$\frac{dn_3}{dt} = -k_{+2}n_2n_3 + k_{-2}n_4 - k_{+6}n_3n_5 + k_{-6}n_9 - k_{+7}n_3n_8 + k_{-7}n_{10}, \quad (3)$$

$$\frac{dn_4}{dt} = k_{+2}n_2n_3 - k_{-2}n_4 - k_{+7}n_4n_8 + k_{-7}n_{13,} \quad (4)$$

$$\frac{dn_5}{dt} = -k_{+3}n_2n_5 + k_{-3}n_6 - k_{+6}n_3n_5 + k_{-6}n_9, \quad (5)$$

$$\frac{dn_6}{dt} = k_{+3}n_2n_5 - k_{-3}n_6 - k_{+4}n_6, \quad (6)$$

$$\frac{dn_7}{dt} = k_{+4}n_6 - k_{+5}n_7, \quad (7)$$

$$\frac{dn_8}{dt} = k_{+5}n_7 - k_{+7}n_8(n_3 + n_4) + k_{-7}(n_{10} + n_{13}), \quad (8)$$

$$\frac{dn_9}{dt} = k_{+6}n_3n_5 - k_{-6}n_9 - k_{+2}n_2n_9 + k_{-2}n_{11} - k_{+3}n_2n_9 + k_{-3}n_{12}, \quad (9)$$

$$\frac{dn_{10}}{dt} = k_{+7}n_8n_3 - k_{-7}n_{10} - k_{+2}n_2n_{10} + k_{-2}n_{13} + k_{+5}n_{14}, \quad (10)$$

$$\frac{dn_{11}}{dt} = k_{+2}n_2n_9 - k_{-2}n_{11}, \quad (11)$$

$$\frac{dn_{12}}{dt} = k_{+3}n_2n_9 - k_{-3}n_{12} - k_{+4}n_{12}, \quad (12)$$

$$\frac{dn_{13}}{dt} = k_{+7}n_4n_8 - k_{-7}n_{13} + k_{+2}n_2n_{10} - k_{-2}n_{13}, \text{ and} \quad (13)$$

$$\frac{dn_{14}}{dt} = -k_{+5}n_{14} + k_{+4}n_{12}. \quad (14)$$

We note that in the previous set of equations $n_j \equiv n_j(t)$ for $j = 1, \cdots, 14$. Rate constants corresponding to phosphorylation events ($k_{+1}$, $k_{+4}$) or molecular dissociation events

($k_{-2}$, $k_{-3}$, $k_{+5}$, $k_{-6}$, $k_{-7}$) have dimensions of inverse time, and thus, units of $s^{-1}$. Rate constants corresponding to molecular association events ($k_{+2}$, $k_{+3}$, $k_{+6}$, $k_{+7}$) have dimensions of inverse concentration and time, and thus, units of $\mu M^{-1}s^{-1}$.

We have assumed that at time $t = 0$, the initial time for the system under consideration, only $F$, $S$ and $P$ are present. Thus, the initial concentrations for all other chemical species vanish. Given the timescales to be studied in the experimental model, the mathematical model does not include protein synthesis, degradation or trafficking of any of the molecular species and hence, the total number of molecules of $F$, $S$ and $P$ are constant in time. We can write down conservation expressions for the total concentration of $F$ ($n_F$), $S$ ($n_S$) and $P$ ($n_P$), since we assume the experimental volume of the system does not change with time. We, therefore, can write

$$n_F = n_1 + n_2 + n_4 + n_6 + n_7 + n_{11} + n_{12} + n_{13} + n_{14}, \tag{15}$$

$$n_S = n_3 + n_4 + n_9 + n_{10} + n_{11} + n_{12} + n_{13} + n_{14}, \tag{16}$$

$$n_P = n_5 + n_6 + n_7 + n_8 + n_9 + n_{10} + n_{11} + n_{12} + n_{13} + n_{14}. \tag{17}$$

We note that in the previous set of constraints $n_j \equiv n_j(t)$ for $j = 1, \cdots, 14$. The above equations hold since the total concentration of a molecule at any time *t* is the sum of the concentrations of all species containing this molecule at that time point.

It is of interest to see which ternary complex species will prevail in the system for sufficiently late times, and thus, we have analysed the steady states of the mathematical model. Steady states can be found by setting the right hand sides of the differential equations to zero and simultaneously solving the resulting equations for the different molecular species in the system. In this case, due to the complexity of the equations, we cannot find explicit expressions for the species at steady state but we find the following implicit equations for a stable steady state solution (denoted by $n^*$)

$$n_1^* = n_5^* = n_6^* = n_7^* = n_9^* = n_{11}^* = n_{12}^* = n_{14}^* = 0, \tag{18}$$

$n_2^* \neq 0,$ (19)

$n_3^* \neq 0,$ (20)

$n_8^* \neq 0,$

$$n_4^* = \frac{k_{+2} n_2^* n_3^*}{k_{-2}}, \quad (21)$$

$$n_{10}^* = \frac{k_{+7} n_3^* n_8^*}{k_{-7}}, \quad (22)$$

$$n_{13}^* = \frac{k_{+2} k_{+7} n_2^* n_3^* n_8^*}{k_{-2} k_{-7}}. \quad (23)$$

Constraints (15)-(17), together with equations (18)-(23) provide a set of implicit polynomial equations for $n_4^*$, $n_{10}^*$ and $n_{13}^*$. It is interesting to observe that the stable steady state defined by the previous equations only provides a non-vanishing value for the ternary complex $pF \cdot S \cdot pP$ (or $n_{13}^* \neq 0$), and that the other ternary complexes, $pF \cdot S \cdot P$, $pF \cdot P \cdot S$ and $pF \cdot pP \cdot S$, are such that $n_{11}^* = n_{12}^* = n_{14}^* = 0$. This is in agreement with the experimental results presented in this manuscript.

The parameters $k_{+2}$, $k_{+3}$, $k_{+6}$, $k_{+7}$ μM.s$^{-1}$ were fixed by the experimental $k_d$ values where, $k_{d,2}$ = 25.1 μM, $k_{d,3}$ = 0.223 μM, $k_{d,6}$ = 1.16 μM, $k_{d,7}$ = 0.48 μM.

$k_{+1} = k_{+4} = 10^0 \mathrm{s}^{-1}$;$k_{-2} = k_{-3} = k_{+5} = k_{-6} = k_{-7} = 10^{-1} \mathrm{s}^{-1}$, and the molecular association rates $k_{+2}$, $k_{+3}$, $k_{+6}$ and $k_{+7}$ with units of $\mu\mathrm{M}^{-1}\mathrm{s}^{-1}$ are fixed by the experimental $k_d$ values where, $k_{d,2} = 25.1$ μM, $k_{d,3} = 0.223$ μM, $k_{d,6} = 1.16$ μM and $k_{d,7} = 0.48$ μM.

### Reaction Figure:

$$F \xrightarrow{k_{+1}} pF$$

$$pF + S \underset{k_{-2}}{\overset{k_{+2}}{\rightleftarrows}} pF \cdot S$$

$$pF + P \underset{k_{-3}}{\overset{k_{+3}}{\rightleftarrows}} pF \cdot P \xrightarrow{k_{+4}} pF \cdot pP \xrightarrow{k_{+5}} pF + pP$$

$$S + P \underset{k_{-6}}{\overset{k_{+6}}{\rightleftarrows}} S \cdot P$$

$$S + pP \underset{k_{-7}}{\overset{k_{+7}}{\rightleftarrows}} S \cdot pP$$

$$\begin{array}{ccccc}
pF \cdot S + pP & \underset{k_{-7}}{\overset{k_{+7}}{\rightleftarrows}} & \color{red}{pF \cdot S \cdot pP} & \underset{k_{+2}}{\overset{k_{-2}}{\rightleftarrows}} & pF + S \cdot pP \\
 & & & & \uparrow k_{+5} \\
 & & & & pF \cdot pP \cdot S \\
 & & & & \uparrow k_{+4} \\
pF \cdot S \cdot P & \underset{k_{+2}}{\overset{k_{-2}}{\rightleftarrows}} & pF + S \cdot P & \underset{k_{-3}}{\overset{k_{+3}}{\rightleftarrows}} & pF \cdot P \cdot S
\end{array}$$
